# Quantification of cellular distribution as Poisson process in 3D matrix using a multiview light-sheet microscope

**DOI:** 10.1101/194571

**Authors:** Warren Colomb, Matthew Osmond, Charles Durfee, Melissa D. Krebs, Susanta K. Sarkar

## Abstract

The absence of quantitative *in vitro* cell-extracellular matrix models represents an important bottleneck for basic research and human health. Randomness of cellular distributions provides an opportunity for the development of a quantitative *in vitro* model. However, quantification of the randomness of random cell distributions is still lacking. In this paper, we have imaged cellular distributions in an alginate matrix using a multiview light-sheet microscope and developed quantification metrics of randomness by modeling it as a Poisson process, a process that has constant probability of occurring in space or time. Our light-sheet microscope can image more than 5 mm thick optically clear samples with 2.9 ±0.4 *μm* depth-resolution. We applied our method to image fluorescently labeled human mesenchymal stem cells (hMSCs) embedded in an alginate matrix. Simulated randomness agrees well with the experiments. Quantification of distributions and validation by simulations will enable quantitative study of cell-matrix interactions in tissue models.

## Introduction

The lack of three-dimensional (3D) *in vitro* models is the missing link in drug discovery^1^ and transport studies^2^. While quantitative analyses of 3D extracellular matrix remodeling by cells have been reported^3^,^4^, the randomness of cell locations in matrices is an unrealized opportunity for quantitative *in vitro* model development for studying the interactions of cells with the extracellular matrices. Randomness is in inherent in biological systems^5^ and plays important roles in diverse biological processes including stem cell proliferation^6^, biochemical reactions^7^, cell signaling^8^, circadian clocks^9^, and gene expression^10^. Similarly, the random distribution of cells in 3D biopolymer matrices that can serve as tissue models is often as random as chocolate chips in cookies. Such randomness can be conveniently modeled and quantified as Poisson process^11^, and a deviation or heterogeneity from an ideal Poisson process can have significance in biology^6^. Quantification of this randomness and cellular tracking in 3D matrices such as hydrogels would require imaging of thick 3D samples. Such an imaging ability may help better define the cell-ECM interactions in these thicker samples. A microscope that can track cells embedded in a several millimeters thick 3D matrix with resolution on the micron scale would be appropriate for diverse studies on cell-ECM interactions. However, out of focus absorption and scattering of the excitation and emission wavelengths can limit 3D imaging. In this regard, the light-sheet microscope (LSM) has proved very useful for a wide range of specimens^12^-^23^ including whole zebrafish^24^, *Drosophila melanogaster*^25^, whole mouse brain^18^,^26^-^28^, and even an entire mouse^29^. LSM involves illuminating only a section of the sample by using a thin light-sheet either created by scanning a focused spot or by using cylindrical lenses. Because of the excitation is limited to a thin layer, LSM enables improved contrast and decreased photobleaching compared to the whole sample illumination. In addition, specimens can be scanned relatively quickly with LSM, enabling dynamic imaging of biological processes on a cellular level. The imaging resolution (nm – few μm) and depths (∼100 μm – few mm) of LSM can vary depending on the method of creating the light-sheet and the optical clarity of the sample, which can be enhanced by chemical cocktails such as CLARITY^30^, Scale^31^, and uDISCO^29^. For significant statistics of cell distributions, however, imaging thicker samples with subcellular resolution of a few micron is more desirable.

In this paper, we present a study on quantifying cellular distributions within a 5mm thick 3D matrix contained in a cuvette by using a lens-based multiview LSM with μ3 μm resolution. Multiview LSM with two detection arms allows rapid imaging with high fidelity because of the fixed geometric arrangement of the detection system^32^,^33^. To detect more emission and thus, to increase the fidelity of detection, we imaged the same layer from two sides of the sample using two 0.5 NA objectives (Fig. 1). To detect more emission and thus, to increase the fidelity of detection, we imaged the same layer from two sides of the sample using two 0.5 NA objectives with 1.1 cm working distances (Fig. 1). The long working distance objective allows us to image cells deep (∼14 mm) into the alginate matrix. We experimentally determined the z-axis resolution using fluorescent beads in agarose gels. On one end, there are light-sheet microscopes with better resolutions, which can only image hundreds of microns at most^34^-^41^. On the other end, some light-sheet microscopes can image centimeter thick samples, but with decreased z-resolution, often tens of microns^29^,^31^,^42^-^44^. These microscopes have largely focused on embryonic development (high resolution) or large scale organ and neuronal connections (decreased resolution). In the middle range, imaging mm-thick samples with micron resolution has also been reported^45^.Therefore, our light-sheet microscope fills the niche area of imaging 5 mm or more thick samples with resolutions suitable for cellular tracking within a 3D matrix. We quantified the randomness of the fluorescent beads and cells by calculating distributions of pairwise distance between locations and validated our approach by modeling randomness as a Poisson process to explain our experimental distributions.

**Fig. 1.**
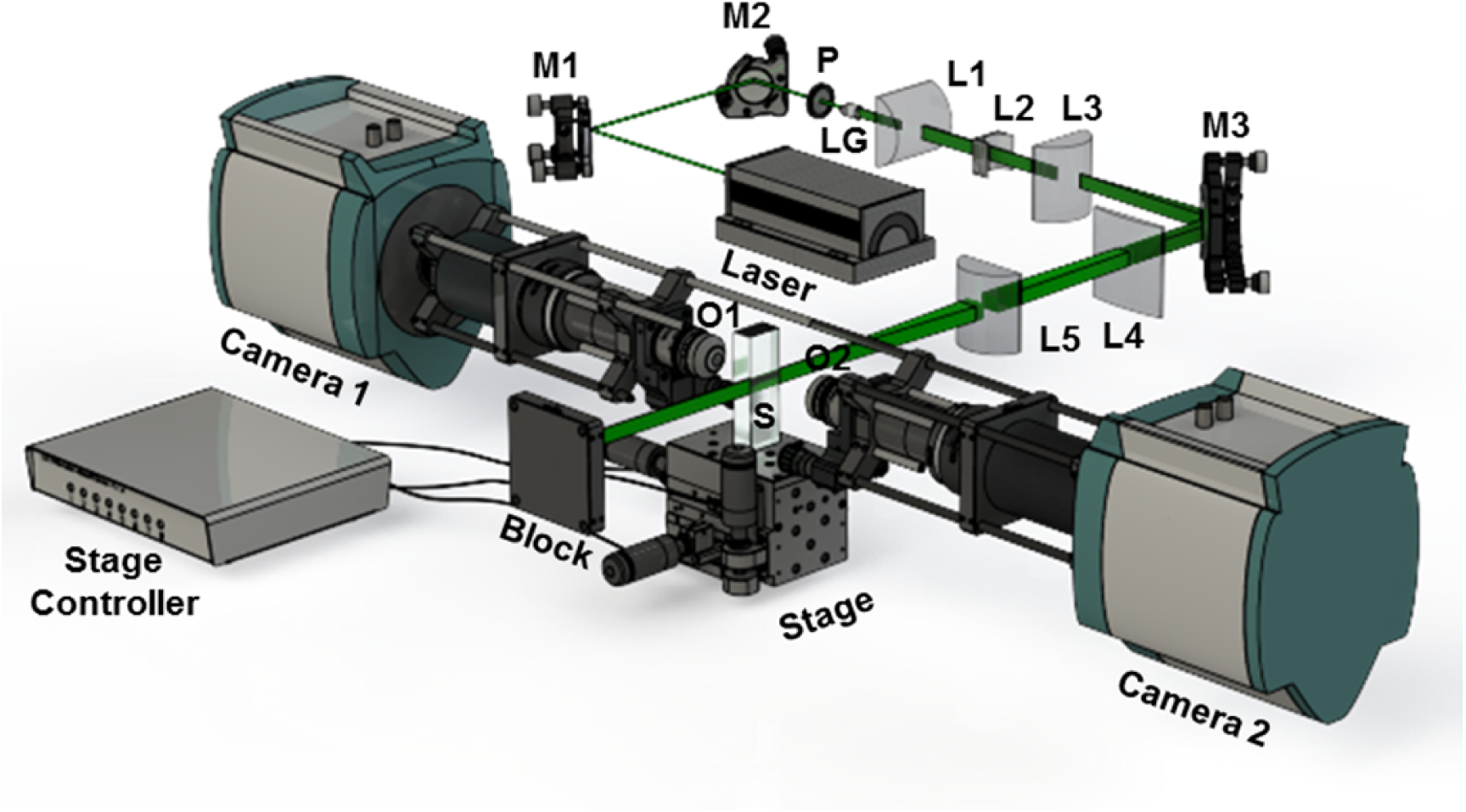
Schematics of the light-sheet microscope. The laser is first expanded into a sheet using a combination of two steering mirrors (M1 and M2), a laser line generator (LG), and three cylindrical lenses (L1, L2, and L3). After reflection from the mirror M3, the laser is compressed horizontally using two cylindrical lenses (L4 and L5) before illuminating the 3D sample (S). Fluorescence from the sample is collected perpendicularly with two 50X long working distance objectives (O1 and O2) in combination with filters and two EMCCD cameras. The detection arms define the z-axis.

## MATERIALS AND METHODS

### Building the multiview light-sheet microscope

We used a commercially available ∼1W 532 nm CW laser as the illumination source. A variable ND filter (Thorlabs, NDC-50C-2M) was used to tune the laser intensity and a polarizer (Thorlabs, WP25M-U-B) was used to polarize the beam if needed. The Gaussian beam from the laser source was converted into a line by using a laser line generator (Edmund Optics, Catalog#43-473, 30μ full fan angle laser line generator lens). The laser line was then collimated to ∼1 cm high beam with a cylindrical lens (Thorlabs, LJ1728L1-A). This vertical beam was expanded horizontally using two cylindrical lenses (Thorlabs, LK1426L1-A and LJ1703L1-A) to create a box-shaped laser profile with dimensions ∼1 cm x 1 cm. The light was then compressed horizontally 6X to a width ∼1 mm using two cylindrical lenses (Thorlabs, LJ1996L1-A and LJ1728L1-A) to create a nominally collimated sheet of light propagating through the sample under study. The light-sheet excites the fluorescent particles embedded in the clear and thick sample within the cuvette. Since the samples contained mostly water, the cuvette was placed in a water chamber to match refractive indices so that the optical thickness to the focal plane of the detection objectives remained constant. We attached the cuvette to a 3D mechanical stage (Newport UlTRAalign 562-XYZ stage with NanoPZ PZA12 actuators) whose micro-step was ∼10 nm (vendor quoted) and ∼13.2±0.2 nm (experimentally measured). Resolution of the linear motion of the stage was measured by tracking particles as the stage was moved in one step increments. The emission from the fluorescent particles was collected from both sides in the direction perpendicular to the excitation direction using a long working distance objective (Olympus, LMPLFLN50X, M PLAN FL 50X objective, NA 0.5, working distance 10.6 mm). The collected emission was optically filtered (Thorlabs 533/17 in combination with Semrock 532U-25) and detected with two separate EMCCD cameras (iXon Ultra 897). The size of a field-of-view was 163 *μm*×163 *μm*. The plane of the light-sheet was called the *xy*-plane, whereas the direction of the detection axis was called the *z*-direction. The light-sheet and the detection arms were fixed, only the sample was moved using the 3D mechanical stage.

### 3D thick sample preparation

Fluorescent beads in agarose

As a reference, we prepared a 5 mm thick sample by embedding fluorescent beads in agarose in a cuvette. We made a 0.5% w/v agarose gel by mixing 100 mg low melting point agarose (Bio-Rad, PCR Low Melt Agarose Cat# 161-3113) with 20 ml ultrapure DI water in a beaker. The mixture was microwaved with a 1200 W microwave for 30 s, swirled to avoid boil over, and then microwaved for another 15 s. The agarose was allowed to cool for approximately 2 min until it was at a safe handling temperature. We then pipetted 5 ml into a test tube to which we added 1 μl of fluorescent bead stock (Bangs Laboratories, FS04F – 1.01μm – PS 525,565 – Envy Green – 1% Solids, sonicated for ∼1 min). We mixed the sample using a micropipette tip for ∼15 s. The mixture was then pipetted into the 5 mm thick cuvette (Sterna Cells, 3-G-5) and covered to cool for ∼1 h. The cuvette was then sealed with a cap and parafilm to keep air out, and to keep the agarose mixture from drying. The outside of the cuvette was then cleaned thoroughly with ethanol and optics tissue paper to remove any debris.

hMSCs in alginate

Human mesenchymal stem cells (hMSCs, from Texas A&M Health Science Center Institute for Regenerative Medicine) were stained with CellTracker Red CMTPX dye (ThermoScientific) according to the manufacturer’s instructions. hMSCs were removed from the plate using 0.05% trypsin for 1 min where they were transferred to a 15 mL conical tube, to be counted and centrifuged to remove excess trypsin and media. The cell pellet was suspended to a concentration of 5×10^5^ cells/mL in PBS 1X solution. 2 mL of a 2 w/v% RGD-alginate solution was made in PBS 1X solution. RGD-alginate was synthesized and purified using a previously publish protocol^46^. 16 μL of 210 mg/mL CaSO_4_ slurry was mixed with 144 μL of the cell suspension containing hMSCs. The RGD-alginate and cell suspension were then mixed between two 3 mL luer syringes for 10 seconds and injected into a cuvette (Sterna Cells, 3-G-5) using a 22-gauge needle. The cuvette was sealed with a cap and parafilm to keep from drying out. The outside of the cuvette was then cleaned thoroughly with ethanol and optics tissue paper.

### Image stack acquisition and processing

A combination of Andor’s imaging software iQ3 and LabVIEW was used to record images with the two cameras as the sample stage moves through ∼1.5-3.5 μm steps in *z*-direction depending on the desired precision. In brief, LabVIEW triggered the Andor EMCCD cameras to start imaging. At the end of image acquisition, the cameras sent a signal to a control program written in LabVIEW which then sent commands to the stage actuators (Newport, PZA12 PZ-SB) to move to the next position. This process was repeated until the entire 3D sample was imaged. The image stack was exported as a multipage TIFF and analyzed using ImageJ plugins and custom MATLAB code. In ImageJ, a 3D Gaussian filter was applied to decrease image noise. Additionally, we used a focus–weighting function to reduce out-of-focus background^47^ by assigning lower weights to out-of-focus intensities than in-focus intensities. The resulting images were smoothed by a user defined amount (generally a 3x3x3 pixel box), and then plotted in 3D for visual analysis with a “black level” set at the peak of the intensity histogram. This allows the high intensity areas to be seen, without the dark areas obscuring them, an issue not present in 2D imaging. We used freely available ImageJ plugins for tracking, MOSAIC and TrackMate, to track fluorescent beads or cells in the processed images. Even though the beads and cells were not moving and were fixed in positions, the tracking program was still useful to find the locations.

### Characterization of the microscope specifications

We quantified the z-resolution by tracking fluorescent beads in agarose. We used two different tracking programs for tracking, MOSAIC and TrackMate in ImageJ. To measure the z-resolution, we used MOSAIC because it provides position as well as intensity moments (zeroth-fourth). The background-free image stack was loaded into the MOSAIC tracking software with the following parameters: Radius 5 (pixels), Cutoff 0, Per/Abs 0.1, Link Range 2 (frames), Link Distance 2 (pixels), Motion Brownian. The resulting tracks were then exported to a text file for analysis in MatLab. The zeroth order intensity (the integrated intensity within the tracking pixel box) was then fitted with a Gaussian function to determine the width of the distribution which was used as our axial resolution metric. Using this metric, the z-resolution is 2.9 ± 0.37μm. The x-y resolution is ∼220 nm.

### Modeling and simulating Poisson process

We analyzed the pairwise distance distributions in 3D (Fig. 4c and 4d) between each pair of beads. For example, if one had 3 particles, the first particle would have two pairwise distances associated with it (1→2, 1→3), the second would have one (2→3), and the third would have already been counted (2→3, 1→3). An image was simulated having the same size as the experimental 3D images (∼163 μm x 163 μm x 5000 μm). For simulation, the image volume was divided into a 3D grid with a voxel size of 0.32 μm x 0.32 μm x 3.3 μm. A random number between 0 and 1 was generated for each grid and filled with particles with a constant probability, *P*, determined from the experimental image according to the equation, *P* =*N* / (*x*(*pixel*)\**y*(*pixel*)\**z*(*pixel*)), where *N* is the number of particles found in a 3D image and *x*(*pixel*), *y*(*pixel*), and *z*(*pixel*) are the number of pixels along each axis. If the value at each voxel (3D pixel) is below *P*, then we considered that a particle was at the voxel. Every other voxel was assumed to be not populated. The pairwise distance distributions for simulated 3D images were analyzed and compared with the experimental distributions.

## Results and discussion

The experimental scheme for the light-sheet microscope is shown in Fig. 1 that combines three microscopy features, i.e., light-sheet illumination using cylindrical lenses^43^, structured illumination using a spatial light modulator^15^, and two detection arms^32^. A thin sheet of light can be created by multiple approaches including cylindrical lenses^43^, scanned Bessel beam^12^, and scanned Gaussian beam^32^. For cellular tracking in a thick 3D matrix, cylindrical lens-based illumination is appropriate and allows for more than 5 mm thick samples to be imaged with ∼3 μm resolution. We used two detection arms to improve the image fidelity instead of sample rotation to avoid the difficulty of setting and accounting for the rotation axis. To measure the resolution of the microscope, a thick test sample made by mixing fluorescent beads in 0.5% (w/v) agarose in a glass cuvette was used. Both the light-sheet illumination and the detection arms were fixed during the image acquisition. We tracked individual fluorescent beads in FIJI with freely available tracking plugins: MOSAIC and TrackMate. Tracking in MOSAIC provides the locations of the fluorescent beads and the intensity moments up to fourth order within the 5-pixel radius. The zeroth order intensity was fitted to a Gaussian, a*exp(-(*x* - *b*)^2^ / *c*^2^), using MATLAB’s built-in curve fitting function fit(), with the ‘gauss1’ argument using the nonlinear least squares method. The fit parameter *c*, the width of the Gaussian, was used as our axial resolution metric. Fig. 2a shows the intensity profiles of different particles as the sample was moved in the *z*-direction. Fig. 2b shows *R*^2^, a measure of goodness of fit was plotted against the width for many particles. Z-resolution, as determined by the mean width in Fig. 2b, was found to be ∼2.9±0.4μm. Fig. 2c shows an example of the intensity profiles along with the fitted Gaussian. Resolution in the *xy*-plane were limited by the bead size with ∼220 nm standard deviation. The theoretical diffraction limited resolution λ/ 2*NA* yields ∼ 0.6 μm lateral resolution for emission at 600 nm. The lateral field of view is ∼163μm X 163μm with 50X objectives and 16μm pixel size of Andor EMCCD cameras. To image the sample volume ∼163μm X ×163μm X 5000μm shown in Fig. 4a, we needed ∼40 min for image acquisition. For this sample, we averaged four 25 ms exposures per image plane and we acquired a total of 1500 image planes, thus the limiting time factor was the motion of the actuators used.

**Fig. 2.**
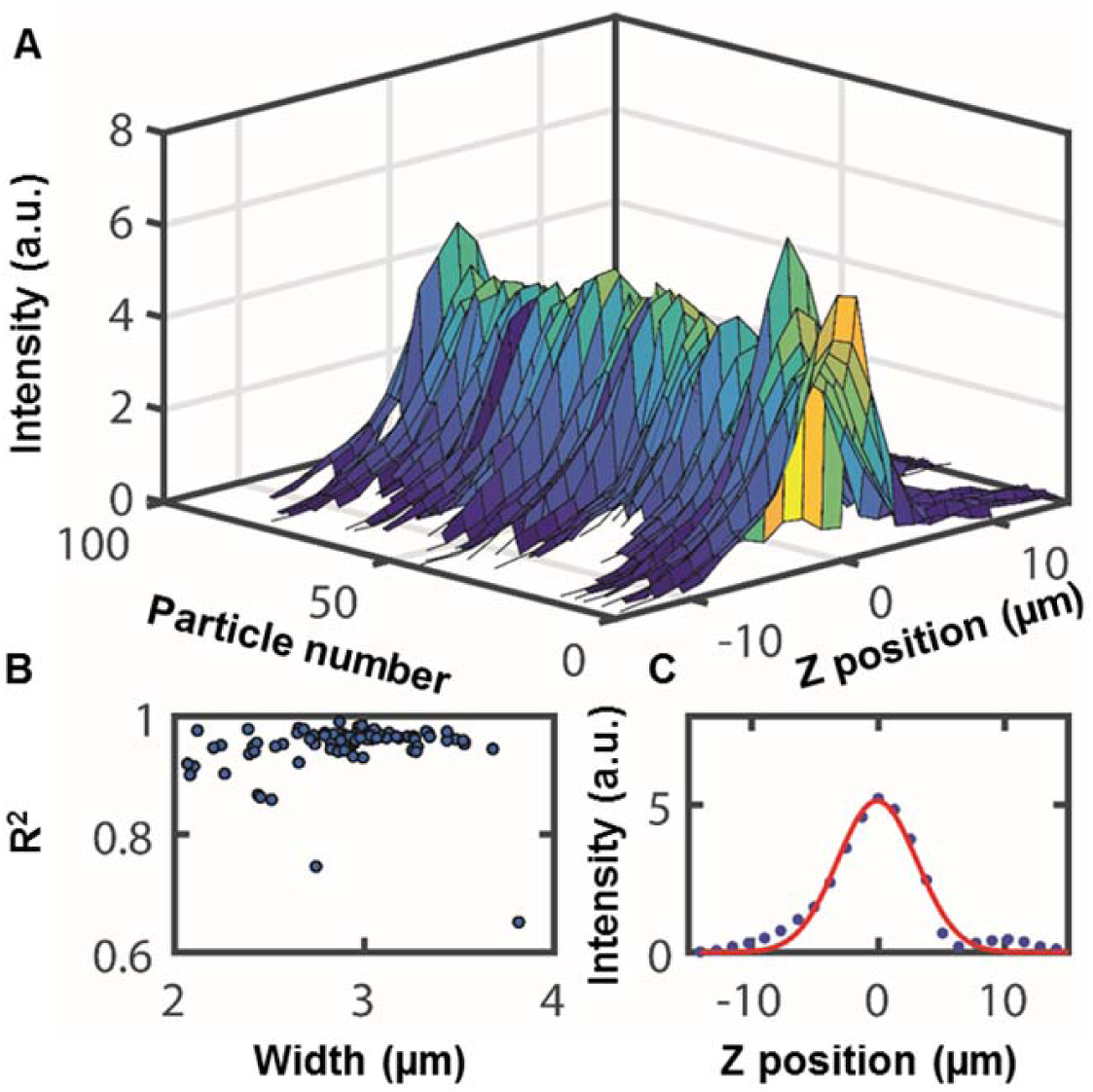
Characterization of the microscope resolution. (A) 3D Plot of the intensity of tracked particles as a function of relative z-position; peak intensity values were set to *z*=0 for all particles. (B) R-Squared of the Gaussian fit for each particle’s intensity track plotted against the FWHM of the fit giving a z-resolution ∼2.9±0.4 μm. (C) An example of intensity profile (blue dots) with its corresponding Gaussian fit (red line).

Both scattering and absorption increase as the sample thickness is increased and become limiting factors for thick sample imaging. Fig.3a shows the composite image of fluorescent beads embedded in alginate gel. The top (a) and bottom (b) 50 slices were then averaged to show more beads in the given field of view. As can be seen, the bottom image (b) has significantly more distortions than the top image (a). Even though the image registration of multiple fields of view in the lateral direction worked well, it is clear that the fluorescent spots show more scattering, less signal, and increased image defects at larger depths. Fig. 3b shows a region of the 3D image to illustrate the typical elongation of the point-spread function along the z-direction, i.e., the optical axis of the detection.

**Fig. 3.**
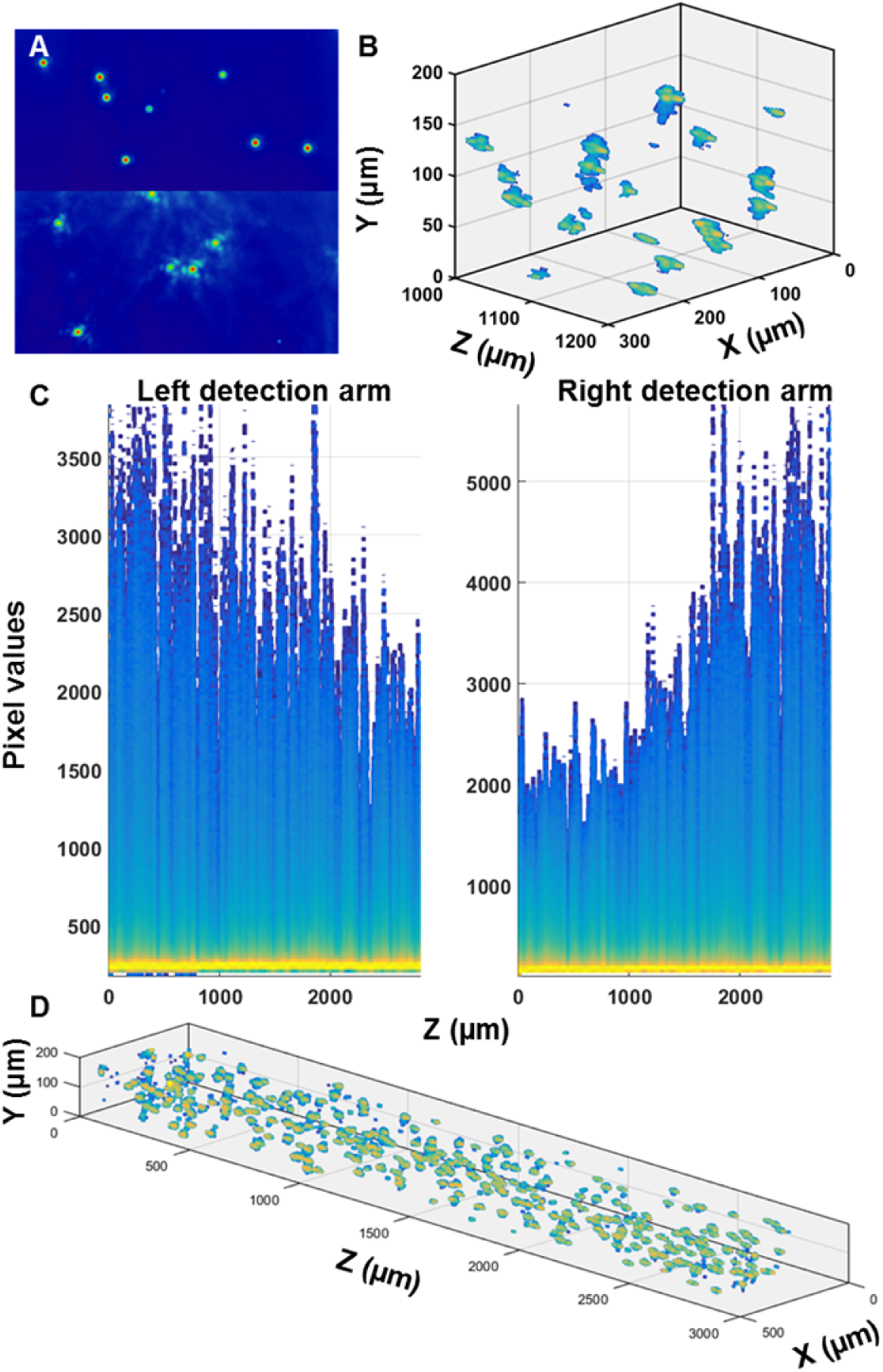
Improved image fidelity by two detection arms. (A) Image of beads in alginate gel taken at a relative depth of zero microns (top) and 5 mm (bottom). (B) Elongated point-spread function along the optical axis of detection. (C) Mesh plot of area-normalized intensity histograms individual image frames. (D) Combined image based on two detection arms.

Fig. 3c shows the intensity histograms at different z-depths detected by two arms and clearly shows that the maximum value as well as the standard deviation of intensity decreases as we imaged deeper in the sample. As expected, the bright spots became less bright and the background became large as the imaging depth was increased. The bright yellow band is the area of highest probability within the histogram and corresponds to the black level of the EMCCD camera. Clearly, imaging thick samples by two detection arms complements the signal quality and was combined to maintain the image quality throughout the sample. Fig. 3d shows the locations of fluorescent beads in agarose gel obtained by combining the images by two detection arms. The background was removed by setting the transparency level to be that of the black level of a normal 2D picture. The composite image was obtained by laterally registering and aligning 5 image stacks each with dimensions163 *μm*X163 *μm*X3320 *μm*. As a result, the ccomposite image had dimensions 433 *μm* X180 *μm* X3840 *μm*, which was then cropped tto 431 *μm* X157*μm* X 2820 *μm* as shown in Fig.3d. Combining images from both image arms homogenized the image intensity as a function of depth, as well as diminished the effects of any abnormality only seen by one camera. For image registration and alignment of the two arms, we used pairwise stitching to align multiple field of views for each detection arm^48^, and then we used descriptor-based registration^49^ to align the 3D images taken by two arms. Stitching with the locations given by sample stage actuators is not precise enough. Instead, a more precise method that calculates Fourier transforms of images, quantifies the cross-correlation, and then globally optimizes the configuration of the combined stitched image was used^48^. To align 3D images from both arms, a global optimization method that is robust with respect to differential movements of the sample was used^49^.

To quantify the quality of alignment, we calculated the distributions of closest points in the two aligned images. The mean distance between closest points is 1.05 *μm*, whereas the median and standard deviation are 0.81 *μm* and 0.67 *μm* respectively. The registration and alignment of two images are below the z-resolution of ∼3 μm as shown in Fig.3. In contrast, the mean, the median, and the standard deviation of distance between closest points without registration are 19.99 *μm*, 19.73 *μm*, and 3.4 *μm* μm respectively; all of which are larger than the z-resolution. Therefore, the registration and alignment are important for multi-view imaging. We quantified random distribution of fluorescent objects by modeling it as a Poisson process which has a constant probability of occurring at every time or spatial step. Examples of Poisson processes are radioactive decay or unbinding of two molecules in the time domain and the distribution of chocolate chips in a cookie or surface deposition of molecules in the spatial domain. There are two consequences of a Poisson process. First, the distributions of times (for time domain Poisson processes) or distances (for spatial Poisson processes) between events are exponential. Second, the distributions of the number of events in user defined blocks of time such as per second and space such as per cubic centimeter can have wide range of shapes including exponential, Poisson distribution, and Gaussian. If there are underlying interactions, the random distributions can have heterogeneities and we need more than one Poisson process to match the data. It should be noted that the Poisson process and distribution are not the same even though Poisson process can lead to Poisson distribution. We quantified the randomness and demonstrated the application of the light-sheet microscope by measuring random distribution of hMSCs suspended in RGD-alginate. In the tissue engineering field, a confocal microscope with an imaging depth limited to a few hundred microns is usually used. However, the ability to optically track cell locations in a thick matrix may be more desirable. While Fig. 4a shows randomly distributed fluorescent beads with size ˜1 *μm* in 0.5% agarose, Fig. 4b shows randomly distributed hMSC cells embedded in alginate matrix. The total number of spots in 3D images were divided by the number of voxels to determine the constant probability of finding a spot. For fluorescent beads (Fig. 4a) the constant probability was 5.66×10^-7^, whereas for cells (Fig. 4b) the probability was 3.9-10^-8^.

**Fig. 4.**
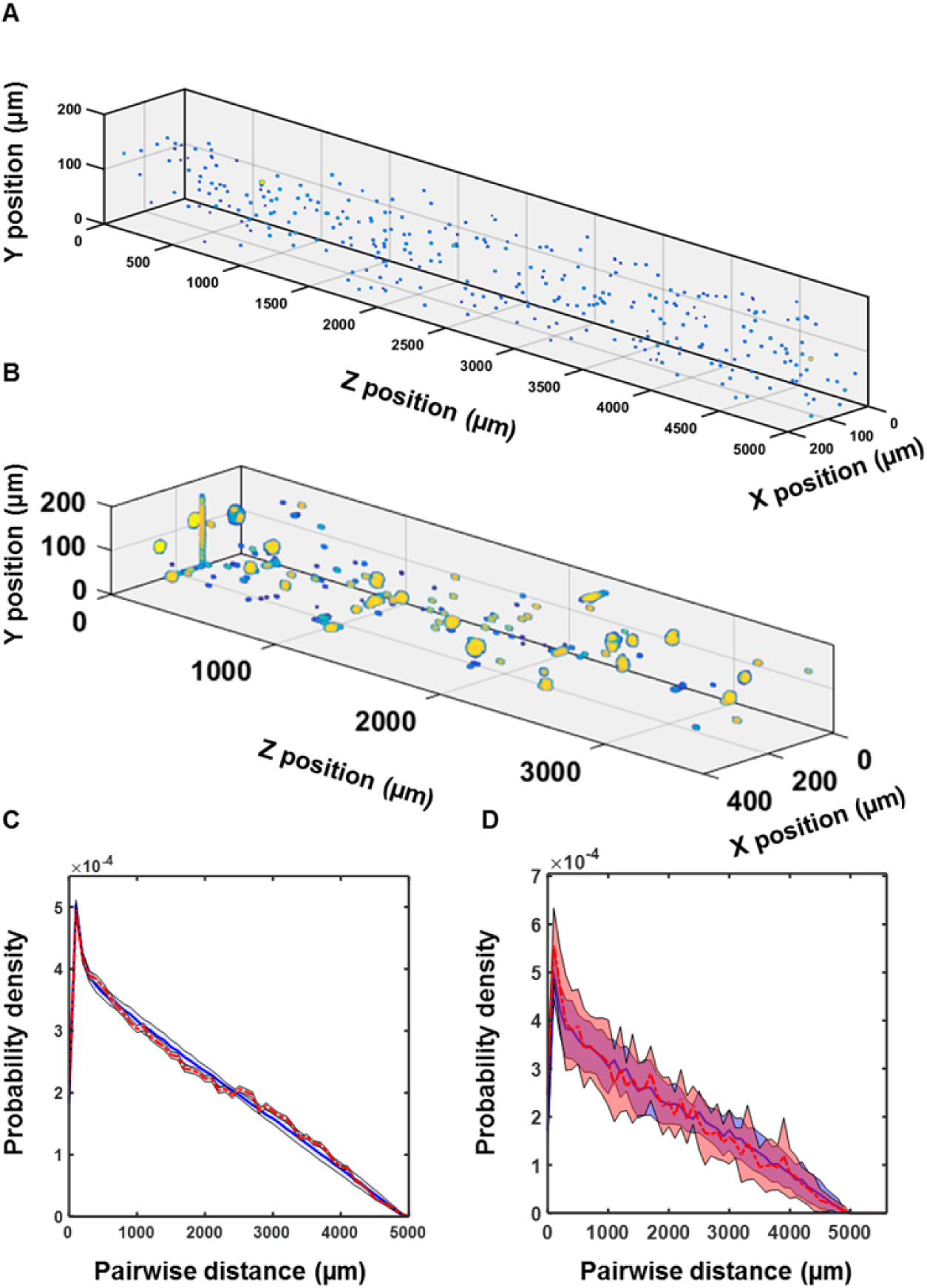
3D imaging of thick samples and quantification of randomness. (A) 3D image data of 1μm sized fluorescent beads in an agarose gel. The background cutoff was set to the peak of the intensity histogram (0.05) so that the beads are visually separated from the background, color is proportional to intensity with blue being lowest and yellow being highest. (B) 3D Light-sheet images of hMSCs labeled with CellTracker Red and embedded in Alginate matrix. The black level is varied to show how background can affect the visualization. (C) Randomness as a Poisson process: experimental (red line) and simulated (blue line) distribution of pairwise distances between beads in Figure 3A. (D) Experimental (red line) and simulated (blue line) distribution of pairwise distances between cells 8 different 3D images like Figure 3B. The error (red shaded area) in experimental distribution is given by the standard deviation of the repeats of experiments. For simulated data, the error (blue shaded area) is represented by the standard deviation of 50 repeats of simulated random population of the image space using the same number of particles found in the image data.

With these experimentally determined probabilities, 3D image data was simulated and analyzed similar to the experimental data. Fig. 4c and 4d show that experimental pair-wise distributions agree well with the pairwise distribution from simulated images with the assumption that the random distributions of beads and cells are Poisson processes. In Fig. 4c, the pairwise distances between beads in 8 field of views from both arms were taken. The mean (red line) and standard deviation (red shaded band around the mean) of 16 pairwise distributions were calculated. Simulated 3D distributions of beads were analyzed exactly the same way to get the mean (blue line) and the standard deviation (blue shaded band) as shown in Fig. 4c. For hMSC cell distributions in alginate matrix, a similar procedure was applied for both the experiments and simulations as shown in Fig. 4d.

## Conclusions

We have developed quantification metrics for random cellular distributions in tissue models using a light-sheet microscope that can image more than 5 mm thick samples with 2.9 ± 0.4 *μm* z-resolution at 600 nm. In comparison to other light-sheet microscopes, our system fills a gap to image more than 5 mm thick samples with a few micron z-resolution, particularly suitable for studying cellular randomness in 3D. We have used our microscope to image hMSC cells embedded in an alginate matrix and modeled the cellular distribution as a Poisson process. Simulation and analysis of cellular locations as Poisson process agrees well with the experimental data. Quantification of randomness will allow the measuring of any deviation from Poisson process and possible heterogeneities arising due to cell-ECM interactions. Our method can be widely applied in diverse studies including cellular motion and invasion through optically clear 3D matrices.

## Acknowledgements

The authors would like to acknowledge Colorado School of Mines Tech Fee funding. We would like to thank Ty Coleman for his work on the schematic as well as John Czerski and Lokender Kumar for their comments and suggestions.

## Author contributions

W.C. and S.K.S designed and performed research, and wrote the manuscript. M.O. and M.D.K. performed cell culture. C.D. suggested the method for quantifying the axial resolution. All authors read and edited the manuscript.

